# Circuit contributions to sensory-driven glutamatergic drive of olfactory bulb mitral and tufted cells during odorant inhalation

**DOI:** 10.1101/2021.09.15.460561

**Authors:** Andrew K. Moran, Thomas P. Eiting, Matt Wachowiak

**Author notes:** **Correspondence:** Matt Wachowiak.

## Abstract

In the mammalian olfactory bulb (OB), mitral/tufted (MT) cells respond to odorant inhalation with diverse temporal patterns that are thought to encode odor information. Much of this diversity is already apparent at the level of glutamatergic input to MT cells, which receive direct, monosynaptic excitatory input from olfactory sensory neurons (OSNs) as well as multisynaptic excitatory drive via glutamatergic interneurons. Both pathways are also subject to modulation by inhibitory circuits in the glomerular layer of the OB. To understand the role of direct OSN input versus postsynaptic OB circuit mechanisms in shaping diverse dynamics of glutamatergic drive to MT cells, we imaged glutamate signaling onto MT cell dendrites in anesthetized mice while blocking multisynaptic excitatory drive with ionotropic glutamate receptor antagonists and blocking presynaptic modulation of glutamate release from OSNs with GABA_B_ receptor antagonists. GABA_B_ receptor blockade increased the magnitude of inhalation-linked glutamate transients onto MT cell apical dendrites without altering their inhalation-linked dynamics, confirming that presynaptic inhibition impacts the gain of OSN inputs to the OB. Surprisingly, blockade of multisynaptic excitation only modestly impacted glutamatergic input to MT cells, causing a slight reduction in the amplitude of inhalation-linked glutamate transients in response to low odorant concentrations and no change in the dynamics of each transient. Postsynaptic blockade also modestly impacted glutamate dynamics over a slower timescale, mainly by reducing adaptation of the glutamate response across multiple inhalations of odorant. These results suggest that direct glutamatergic input from OSNs provides the bulk of excitatory drive to MT cells, and that diversity in the dynamics of this input may be a primary determinant of the temporal diversity in MT cell responses that underlies odor representations at this stage.

## Introduction

In the mammalian olfactory system, the neural representation of olfactory information is inherently dynamic, with respiration and active odor sampling (i.e., sniffing) driving inhalation-linked bursts of activity in olfactory sensory neurons (OSNs) that are passed on to higher-order neurons, including mitral and tufted (MT) cells, the main output neurons of the mammalian olfactory bulb (OB) (Schaefer and Margrie, 2007;Wachowiak, 2011). At the same time, OSNs and MT cells show slower temporal patterning across respiration cycles (Patterson et al., 2013;Eiting and Wachowiak, 2020). Temporal patterning at each of these timescales is hypothesized to play important roles in encoding odor information at the level of the OB and in piriform cortex, a main target of MT cell projections (Schaefer and Margrie, 2007;Stern et al., 2018;Chong et al., 2020). The cellular and circuit mechanisms underlying temporal patterning at the level of MT cell output from the OB remain unclear.

Potential contributions towards the temporal patterning of MT cell activity include diverse patterns of OSN activation (Spors et al., 2006;Short and Wachowiak, 2019), presynaptic inhibition of glutamate release from OSN terminals (McGann et al., 2005;Wachowiak et al., 2005), or neural circuits of the OB. Potential OB circuit mechanisms for shaping MT cell output dynamics have been well-characterized using OB slice preparations, and include modulation of glutamate release from ET cells onto MT cells (Hayar et al., 2004a;De Saint Jan et al., 2009;Gire et al., 2012) as well as feedforward inhibition via several distinct inhibitory interneuron pathways (Murphy et al., 2005;Gire and Schoppa, 2009;Shao et al., 2012). In a recent study using glutamate and Ca^2+^ imaging from the mouse OB in vivo (Moran et al., 2021), we found that glutamatergic signaling onto MT cells showed complex dynamics both within and across inhalations and that the slower temporal patterning was well-correlated with that of MT cell Ca^2+^ signals, highlighting the importance of excitatory pathways in generating MT cell patterning.

Here, we sought to dissect the contribution of OSNs versus second-order OB circuits to generating dynamic glutamatergic signaling onto MT cells. We used *in vivo* pharmacology to examine the contributions of these different sources to the dynamics of MT cell glutamatergic input in anesthetized mice. We directly imaged odorant-evoked glutamate signaling onto MT cells while pharmacologically blocking multisynaptic excitatory drive with ionotropic glutamate receptor antagonists and blocking presynaptic modulation of glutamate release from OSNs with GABA_B_ receptor antagonists. Neither manipulation substantially impacted the inhalation-linked temporal dynamics of glutamate signaling onto MT cells. Furthermore, blocking multisynaptic excitation only weakly reduced the magnitude of glutamatergic excitation, and modestly impacted glutamate dynamics over a slower timescale spanning multiple inhalations of odorant. Overall, these results suggest that direct glutamatergic input from OSNs provides the bulk of excitatory drive to MT cells and that this direct OSN – MT cell pathway may be the primary determinant of inhalation-linked temporal patterning of MT cell activity.

## Results

### Presynaptic inhibition does not impact inhalation-linked dynamics of glutamate signaling

Neurotransmitter release from OSNs can be modulated by presynaptic dopamine (D2) and GABA_B_ receptors, which modulate Ca^2+^ influx into the presynaptic terminal and subsequent release of neurotransmitter (Ennis et al., 2001;Murphy et al., 2005;Wachowiak et al., 2005;Vaaga et al., 2017). GABA_B_-mediated presynaptic inhibition is activated both tonically and via sensory-evoked, intraglomerular circuits that have been proposed to mediate gain control in vivo in a manner independent of inhalation frequency (McGann et al., 2005;Pírez and Wachowiak, 2008;Shao et al., 2009;Vaaga and Westbrook, 2017). Thus we first sought to characterize how GABA_B–_ mediated presynaptic inhibition modulates the gain and dynamics of glutamatergic drive onto MT cells in vivo.

First, we confirmed effective blockade of GABA_B_-mediated presynaptic inhibition by imaging Ca^2+^ influx into OSN terminals in dorsal OB glomeruli using the genetically-encoded calcium reporter GCaMP6f expressed exclusively in OSNs (Wachowiak et al., 2013;Rothermel and Wachowiak, 2014;Short and Wachowiak, 2019) and comparing responses before and after topical application of the GABA_B_ receptor antagonist CGP35348 (1 mM) to the dorsal OB, as done previously (Wachowiak et al., 2005;Brunert et al., 2016). Odorant was presented at suprathreshold concentrations (17-48 ppm; Table 1) to anesthetized mice using an artificial inhalation paradigm and evoked signals imaged from the dorsal OB under epifluorescence. Robust odorant-evoked presynaptic Ca^2+^ signals were apparent in multiple foci representing glomeruli across the dorsal OB (Fig. 1A). Temporal response patterns during 2 Hz inhalation consisted of inhalation-linked transients apparent atop a sustained signal (Fig. 1B). CGP35348 application caused an increase in the odorant-evoked Ca^2+^ signal that was apparent as early as the first inhalation of odorant and which persisted throughout odorant presentation (Fig. 1B). The magnitude of the increase varied across glomeruli and in different mice (Fig. 1C), with a mean increase of 97% across the 5 mice tested (mean ± SD of median CGP35348/Predrug ratio per experiment: 1.97 ± 0.34; p=0.003, t-test on median ratios with ratio = 1 as null hypothesis; df = 4). CGP35348 did not appear to change the dynamics of inhalation-linked GCaMP6f transients (Fig. 1B, bottom). These results confirm previous findings using calcium-sensitive dyes (Wachowiak et al., 2005;Pírez and Wachowiak, 2008).

**Figure 1.**
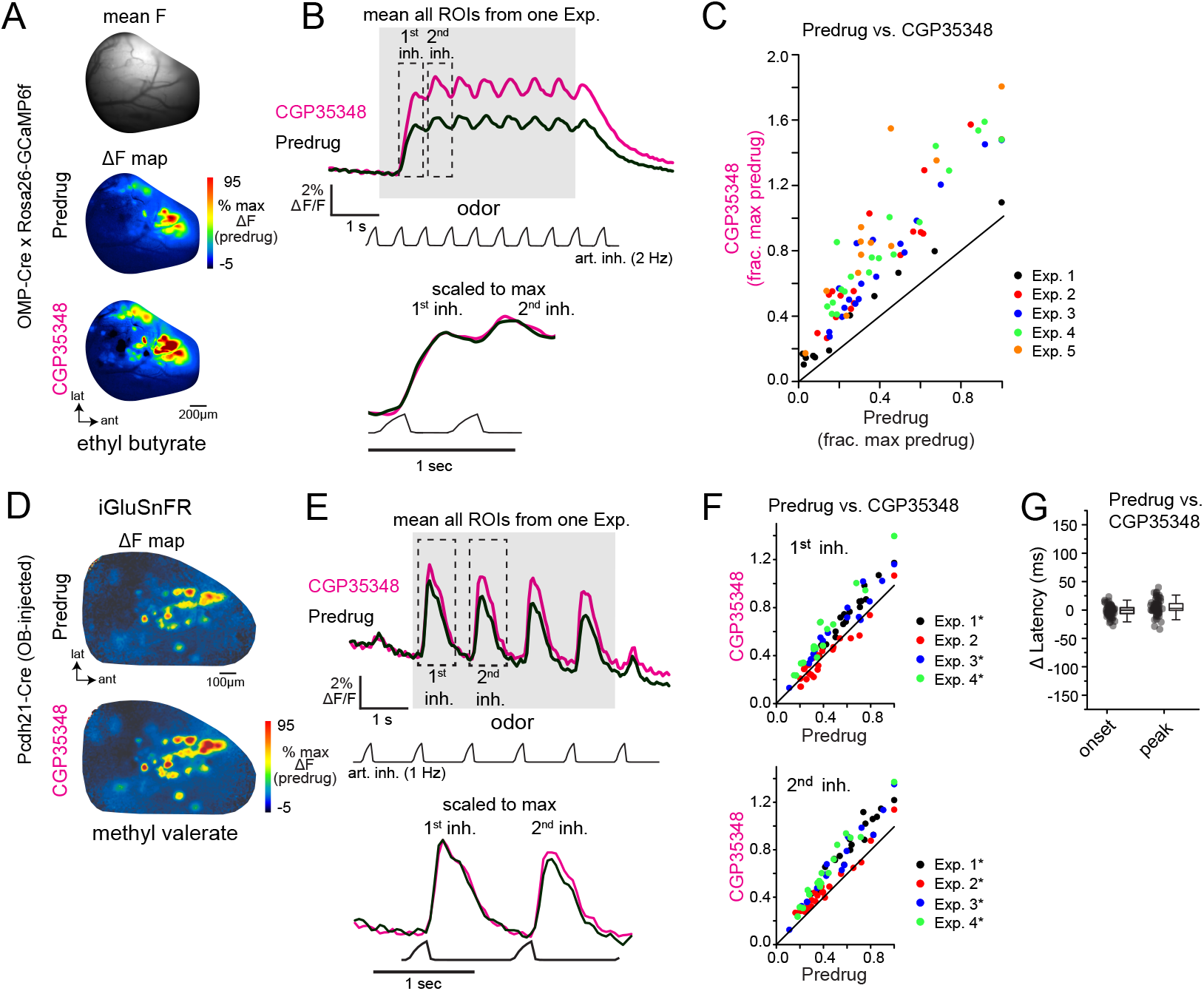
GABA_B_-mediated presynaptic inhibition regulates the strength, but not the timing, of inhalation-driven glutamatergic signaling onto mitral/tufted cells. **A.** Baseline fluorescence (mean F) and ΔF maps (OMP-Cre x Rosa26-GCaMP6f) of calcium responses to methyl valerate before (Predrug) and after application of CGP35348. **B.** Traces showing mean fluorescence of all glomerular ROIs in one OB to odorant presentation (ethyl butyrate) before and after application of CGP35348. Trace showing artificial inhalation pressure is below the response traces (2 Hz). Lower traces show expansion of responses to the first and second inhalation of odorant scaled to the same maximum. **C.** Peak OSN – GCaMP6f responses (ΔF/F) in each measured ROI, normalized to the maximal predrug response in each experiment. Each color is an individual experiment. Summary statistics for each experiment (median CGP35348/Predrug ratio, Wilcoxon Signed Ranks Test, p-value corrected for multiple comparisons): Exp. 1: 1.60, Z = −2.89, p = 0.004, Exp. 2: 1.91, Z = −3.60, p = 3.21 x 10^-4^, Exp 3: 1.76, Z = −3.70, p = 2.14 x 10^-4^, Exp. 4: 2.13, Z = −3.80, p = 1.43 x 10^-4^, Exp. 5: 2.46, Z = −3.02, p = 0.003. **D.** Odorant-evoked ΔF maps showing glutamate signals (SF-iGluSnFR.A184V) on MT cells before and after CGP35348 application. **E.** Mean iGluSnFR fluorescence across all glomerular ROIs in one OB to odorant presentation (ethyl butyrate) before and after CGP35348 application. Odorant was delivered using 1 Hz artificial inhalation. Bottom traces show expanded responses to first two inhalations of odorant, scaled to the peak of the first inhalation response. **F.** Peak MT Cell - iGluSnFR ΔF/F responses (ΔF/F) in each measured ROI before and after CGP35348 application, normalized to the maximal Predrug response in each experiment. Summary statistics, reported as in (C): CGP35348/Predrug (first inhal.) (mean ± SD): Exp. 1: 1.2 ± 0.06, n = 19, p 2.9 x 10^-10^; Exp. 2: 0.97 ± 0.17, p =1.0, n=20; Exp. 3: 1.24 ± 0.15, n=16, p = 2.6 x 10^-5^; Exp 4: 1.4 ± 0.2, n = 14, p = 2.5 x 10^-5^. CGP35348/Predrug (second inhal): Exp. 1: 1.34 ± 0.11, p = 3.0 x 10^-10^, Exp 2: 1.15 ± 0.14, p = 3.9 x 10^-4^, Exp 3: 1.34 ± 0.20, p = 1.7 x 10^-5^, Exp 4: 1.48 ± 0.13, p = 2.1 x 10^-8^. Asterisks indicate Experiments with p < 0.05 after correction for multiple comparisons). **G.** Change in onset and peak latencies of iGluSnFR response to the first inhalation of odorant after application of CGP35348 (shaded magenta, from Predrug). Same mice and glomeruli as in E, F. Each point is a glomerulus; boxes indicate 25th and 75th percentiles, line indicates median, whiskers denote outliers with a coefficient of 1.5. CGP35348 had little to no effect on onset and peak latencies of response to first inhalation (Wilcoxon signed ranks test, medians and p-values per experiment): Δ onset latencies, Exp. 1: −6.3 ms, p = 0.04; Exp. 2: 3.5 ms, p = 0.2; Exp. 3: −3.1 ms, p = 0.016; Exp. 4: 4.9 ms, p = 0.16. Δ peak latencies, Exp 1: 3.3 ms, p = 0.16; Exp. 2: 8.4 ms, p = 8 x 10^-4^; Exp. 3: −2 ms, p = 0.32; Exp. 4: 2.8 ms, p = 0.52.

**Table 1.** Odorants and concentration used. Each row represents a different odorant used in each experimental manipulation. Each column represents dataset in which the pharmacological or stimuli were varied. Final Estimated ppm (by Experiment type) is the estimated vapor concentration of each odorant delivered to the animal, calculated from liquid dilution ratios used, reported vapor pressures, and calibration of the odor delivery device. Final row (‘Figures’) lists figures containing data from each Experiment Type.

| Odorant | Final Estimated ppm<br>(by Experiment Type) |  | Odor<br>Diversity<br>Exp. |
| --- | --- | --- | --- |
|  | High [Odor]<br>0.5 mM NBQX<br>/ 1 mM APV | Low [Odor]<br>2.5 mM NBQX<br>/ 5 mM APV |  |
| 2-Methylbutyric acid |  |  | 0.03 |
| Isovaleric acid |  |  | 0.04 |
| Trans-2-methyl-2-butenal |  |  | 0.7 |
| 2-Methylvaleraldehyde |  |  | 3 |
| Butyl acetate | 17 |  |  |
| Ethyl butyrate | 20 | 1 - 3 | 3 |
| Vinyl butyrate | 3 |  | 2 |
| Methyl valerate | 40 | 2 - 5 | 2 |
| Ethyl tiglate |  | 0.07 - 0.7 | 0.6 |
| Hexyl tiglate |  |  | 1 |
| Isopropyl tiglate |  |  | 5 |
| 2-hexanone | 48 |  |  |
| Cyclohexylamine |  |  | 15 |
| n-Methyl piperidine |  |  | 1 |
| Figures | 1,2,3,4,5 | 3,5 | 6 |

Next, we used the second-generation iGluSnFR, SF-iGluSnFR.A184V, to directly monitor the impact of GABA_B_ receptor blockade on odorant-evoke glutamate signaling onto MT cells. As expected from an earlier characterization using first-generation iGluSnFR (Brunert et al., 2016), CGP35348 led to an increase in the amplitude of inhalation-driven glutamate transients (Fig. 1D, E). Here, in contrast to the GCaMP6f imaging results, we observed a difference in the magnitude of the effect of GABA_B_ receptor blockade on responses to the first and second inhalations of odorant. Overall, CGP35348 caused a significant increase in response to the second inhalation in all 4 mice but increased responses to the first inhalation of odorant in only 3 of 4 mice (Fig. 1F; see Figure 1 legend for statistical tests).

However, GABA_B_ receptor blockade had little to no impact on the inhalation-linked dynamics of glutamatergic signaling onto MT cells. Neither onset latencies nor peak latencies (defined as time to 10% of peak and time to 90% of peak, respectively) of the iGluSnFR transient evoked by the first inhalation of odorant changed substantially after CGP35348 application (Δ onset latency, −0.3 ± 2.6 ms; Δ peak latency, 3.1 ± 4.2 ms; mean ± SD of median values from 4 mice; Fig. 1G). While 2 of 4 mice showed a significant change in onset or peak latency (see Fig. 1G legend), the magnitude of the change in these experiments (<10 ms) was negligible compared to the range of onset and peak latencies seen across different glomeruli in the same preparation, which ranges over a span of 200 − 250 ms (Moran et al., 2021). These results confirm that GABA_B_-mediated inhibition regulates the strength of glutamatergic signaling from presynaptic sources onto MT cells in vivo (McGann et al., 2005;Wachowiak et al., 2005;Brunert et al., 2016), and suggest that presynaptic inhibition contributes little to the inhalation-linked timing of these signals.

To assess the impact of this GABA_B_-mediated modulation of glutamatergic drive on postsynaptic responses of MT cells in vivo, we returned to Ca^2+^ imaging, expressing GCaMP6f in MT cells and comparing odorant-evoked responses before and after CGP35348 application. GCaMP6f signals were imaged from the apical tufts of MT cells across the glomerular layer using two-photon imaging, as done previously (Economo et al., 2016;Short and Wachowiak, 2019) (Fig. 2A, B). CGP35348 had variable effects on MT cell Ca^2+^ responses in different glomeruli, with increased responses in some glomeruli and decreased responses in others (Fig. 2B, C). Suppressive responses, reflected as a decrease in GCaMP6f fluorescence and presumably reflecting inhibition of ongoing MT cell spiking (Economo et al., 2016), persisted after application of CGP35348 (Fig. 2B, C) indicating that MT cell suppression is largely not mediated by presynaptic inhibition of glutamatergic drive from OSNs. Overall, there was no significant net change in response magnitude, either for peak responses to the first inhalation of odorant, or averaged across the duration of odorant presentation (CGP35348/Predrug ratios, first inhalation of odorant (median [quartile range] of medians: 0.75 [0.64 ± 1.05], p = 0.11, one-sample Wilcoxon Signed Rank test, n = 12; odor duration: 0.91 [0.77 ± 1.33], p = 0.85). Thus, increasing the magnitude of excitatory input to the glomerulus by enhanced release of glutamate from OSN terminals does not uniformly translate into increased excitation of MT cells.

**Figure 2.**
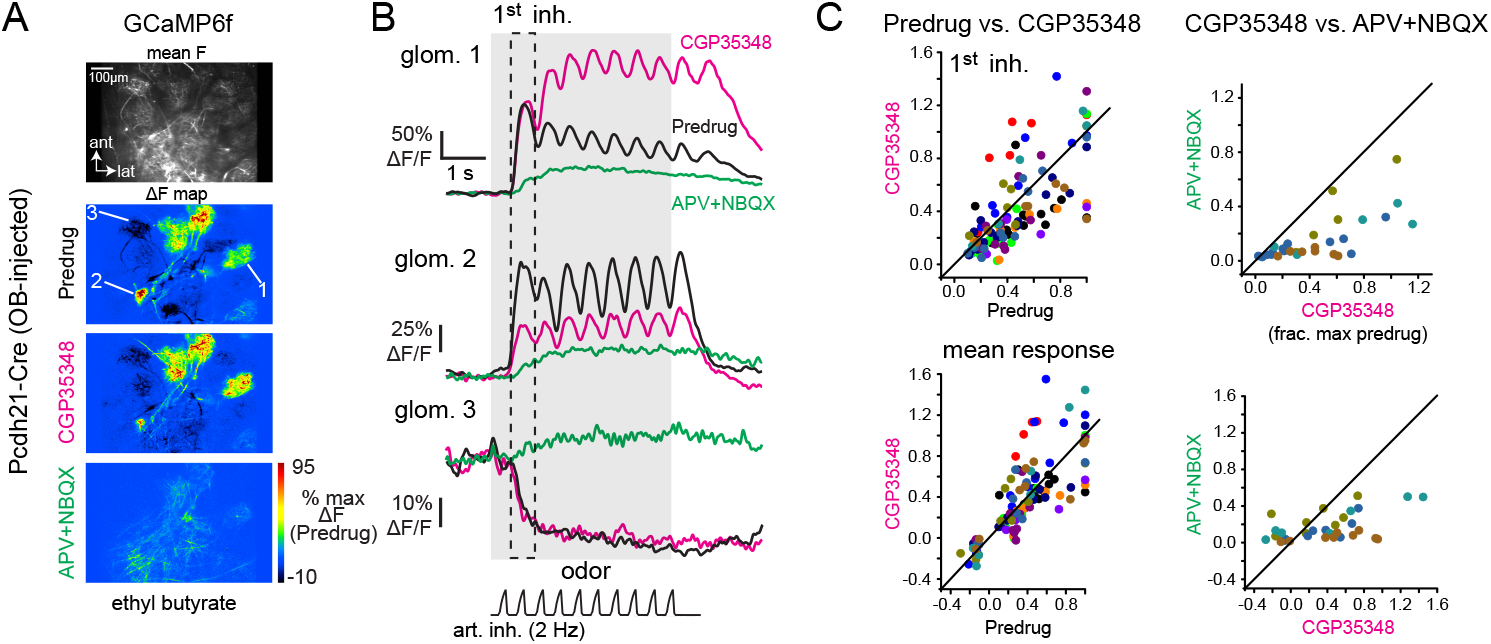
GABA_B_-mediated presynaptic inhibition has a heterogeneous impact on mitral/tufted cell calcium responses. **A.** Baseline fluorescence (mean F) and ΔF maps of odorant-evoked GCaMP6f signals in mitral/tufted cells, imaged with two-photon microscopy; z-plane is in the deep glomerular layer/outer external plexiform layer of the dorsal OB. Maps show responses to the same odorant (ethyl butyrate) before and after application of CGP35348, and after subsequent application of APV+NBQX. Pseudocolor scale normalized to Predrug response levels. APV+NBQX strongly reduced excitatory responses and eliminated suppressive responses. **B.** Response traces for the three glomeruli indicated in (A), (2 are excited, 1 is suppressed) across each condition. **C.** Comparison of MT cell GCaMP6f responses for Predrug versus CGP35348 conditions (12 mice; left plots) and CGP35348 and subsequent APV+NBQX (4 mice; right plots) conditions. Colors indicate individual mice. Top row: peak response amplitudes to the first inhalation of odorant, normalized to the maximum Predrug response in each mouse. Bottom row: mean response across odorant presentation. See Text for CGP35348 summary statistics across mice. For APV+NBQX/CGP35348 statistics, ratios per experiment, reported as medians [quartile range] were: Exp. 1: 0.47 [0.37 – 0.9], n = 6 ROIs; Exp 2: 0.39 [0.25 - 0.52], n = 11; Exp 3: 0.38 [0.19 – 0.57], n = 5; Exp 4: 0.19 [0.08 – 0.27], n = 6; Full duration: Exp 1: 0.66 [0.46 – 1.12], n = 6; Exp. 2: 0.4 [0.31 – 0.51]; Exp 3: 0.37 [0.27 – 0.50], n = 4; Exp 4: 0.12 [0.05 – 0.18], n = 7. ROIs were pooled across the 4 mice for statistical comparison due to the small number of ROIs in each experiment.

### Limited contribution of multisynaptic pathways to odorant-evoked glutamate signaling onto MT cells

Sensory-evoked glutamatergic input onto MT cells can arise from OSNs or from multisynaptic excitation involving dendritic glutamate release from ET cells and, possibly, sTCs (Hayar et al., 2004a;De Saint Jan et al., 2009;Gire et al., 2012;Vaaga and Westbrook, 2016;Sun et al., 2020). In addition, inhibitory circuits can modulate multisynaptic glutamate signaling (Gire and Schoppa, 2009;Shao et al., 2012;Gire et al., 2019). Glomerular glutamate signals may also arise from glutamate release by MT cell dendrites (Isaacson, 1999;Najac et al., 2015), and thus at least partially reflect MT cell excitation itself. To isolate the direct contribution of OSN inputs to MT cell excitation dynamics, we compared evoked glutamate dynamics before and after pharmacological blockade of postsynaptic activity with ionotropic glutamate receptor antagonists (Gurden et al., 2006;Pírez and Wachowiak, 2008;Lecoq et al., 2009). Because iGluSnFRs are insensitive to these antagonists (Marvin et al., 2013;Marvin et al., 2018), we reasoned that this approach would largely prevent OSN-driven excitation of postsynaptic neurons and allow us to image evoked MT cell glutamate signals arising solely from OSN inputs.

To confirm our ability to block ionotropic glutamate receptors with the in vivo drug application protocol, we first continued the GCaMP6f imaging from MT cells in the same mice used for the GABA_B_ blockade experiments, applying a cocktail of APV+NBQX (1 mM/ 0.5 mM) following the initial CGP35348 treatment (n = 4 mice). APV+NBQX sharply reduced odorant-evoked GCaMP6f signals, such that distinct inhalation-linked transients were eliminated in most glomeruli and replaced with a weak tonic glutamate signal (Fig. 2B); this signal might reflect modulation by metabotropic glutamate receptors (Matsumoto et al., 2009;Dong and Ennis, 2014), which we did not attempt to block in these experiments. Excitatory responses to both the first inhalation and over the duration of odorant presentation were reduced to 36 ± 20% (p = 3.5 x 10^-10^, n = 29 ROIs from 4 mice, paired t-test); and 36 ± 15% (p = 1.2 x 10^-6^, n = 23 ROIs, paired Wilcoxon Signed Ranks) respectively, of their baseline values (Figs. 2B, C) (see Figure legend for summary statistics). Suppressive responses were eliminated after APV+NBQX application and were replaced with a slow, tonic increase (Fig. 2B).

We next tested the ability of APV+NBQX to block activation of cholecystokinin (CCK)-expressing sTCs, as these, like ET cells, are strongly driven by monosynaptic OSN input (Isaacson, 1999;Najac et al., 2015). We imaged activation of CCK+ OB neurons in CCK-IRES-Cre mice crossed to a Cre-dependent GCaMP6f reporter line (n = 3 expts., 2 mice). We first blocked GABA_B_-mediated presynaptic inhibition with CGP35348 to further enhance transmitter release from OSNs. As expected, CGP35348 led to a modest increases in the peak CCK+ GCaMP6f response, with median increases of 33, 9 and 24%, respectively, in the three experiments (Fig. 3A - E). Subsequent application of APV+NBQX (1 mM/0.5 mM) had mixed effects in different glomeruli, with responses completely or nearly eliminated in 8 of 14 glomeruli (97 ± 3% median reduction in peak response to 1^st^ inhalation) but only partially reduced in the remaining 6 (38 ± 16% median reduction) (Fig. 3A-E). In these 6 glomeruli, responses to successive inhalations after odorant onset were reduced even further or eliminated (e.g., Fig. 3E). We performed additional experiments using a 5x higher concentration of antagonist (5 mM APV/2.5 mM NBQX) and without preapplication of CGP35348. In these, APV+NBQX completely blocked postsynaptic activation in 11 of 13 glomeruli (from 3 experiments), with strongly reduced responses (66 and 75% reduction) in the remaining two (Fig. 3F).

**Figure 3.**
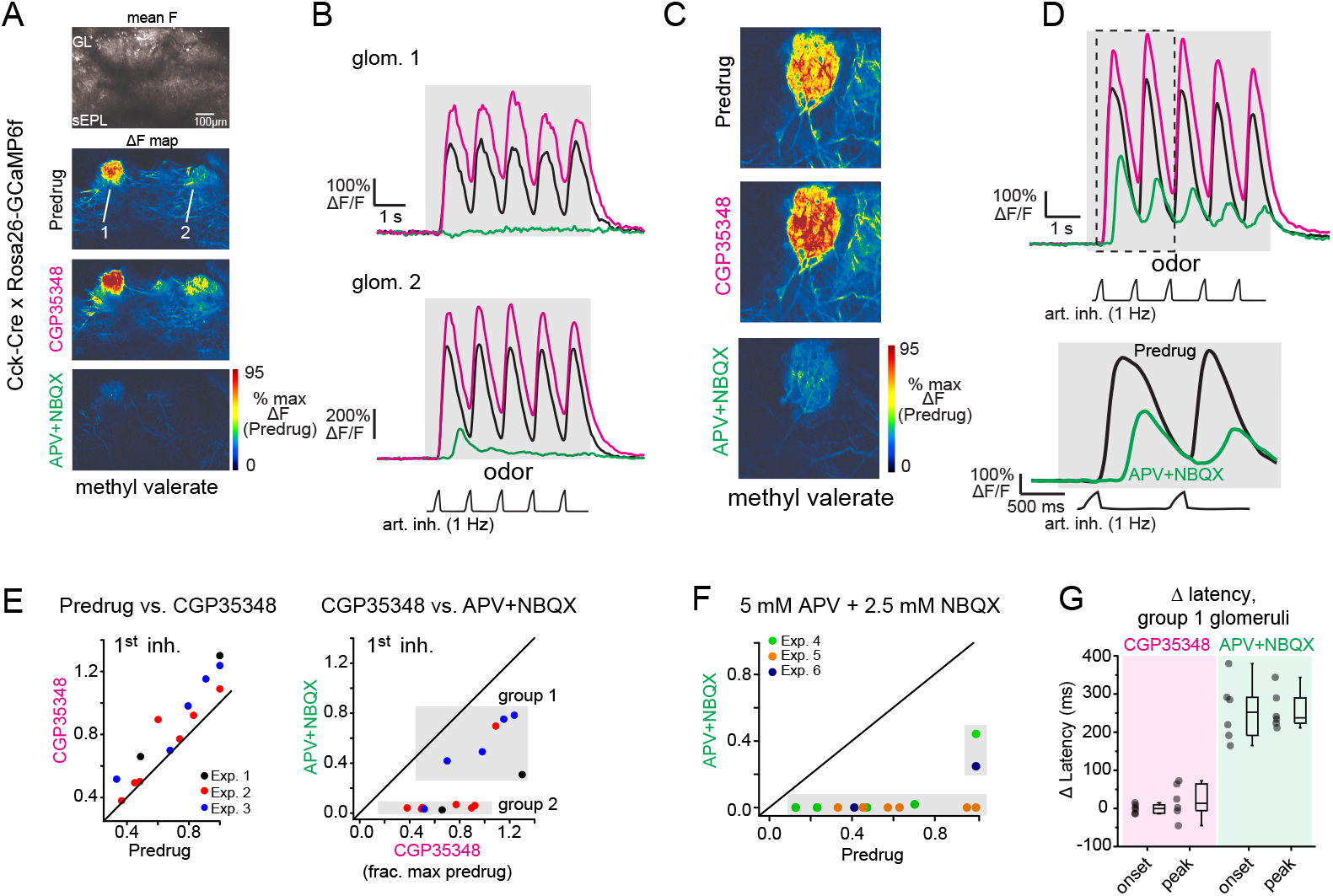
Near-complete blockade of odorant-evoked excitation of CCK+ superficial tufted cells by APV+NBQX. **A.** Odorant response maps of GCaMP6f signals imaged from CCK+ superficial tufted cells, taken across Predrug, CGP35348 and APV+NBQX conditions, scaled to Predrug response levels. **B.** Response traces for the two glomeruli indicated in (A), showing enhanced response amplitudes after application of CGP35348 and either elimination (top) or sharp reduction (bottom) in responses after subsequent application of APV+NBQX. Note that remaining response after APV+NBQX is delayed. **C.** Example of high-zoom imaging of GCaMP6f signals imaged from superficial CCK+ neurons innervating a glomerulus, with response maps in Predrug, CGP35348, and APV+NBQX conditions. **D.** Traces showing odorant-evoked response from glomerular neuropil in (C). CGP35348 increases response amplitudes. APV+NBQX decreases response amplitude, with delay in remaining signal relative to inhalation. Lower traces show expansion of response to first two inhalations of odorant, highlighting delayed GCaMP6f response after APV+NBQX. **E.** Comparison of CCK+ GCaMP6f responses across CGP35348 and subsequent APV+NBQX conditions, imaged from 14 glomeruli in 3 OBs (2 mice). Scatter plots show peak response to first inhalation of odorant in Predrug compared to CGP35348 condition (left) or CGP35348 and APV+NBQX conditions (right). Shaded regions in APV+NBQX vs. CGP35348 plot (upper right) indicate two groups of glomeruli defined by the magnitude of response reduction by APV+NBQX (see Text). **F.** Comparison of CCK+ GCaMP6f responses before (Predrug) and after higher concentrations of APV+NBQX (5 mM/2.5 mM), and without prior application of CGP35348. Separate experiments from (E). Responses are fully blocked in 11 of 13 glomeruli, (3 experiments). **G.** Change in onset and peak latencies of CCK+ GCaMP6f response to the first inhalation of odorant after application of CGP35348 and APV+NBQX, for the 6 group 1 glomeruli shown in (E). Response latencies for group 1 glomeruli are unchanged by CGP35348 application (left), but increase significantly after APV+NBQX application (right). Summary statistics, mean ± SD, paired t-test: CGP35348 vs. Predrug Δ onset latency: −0.5 ± 12 ms, p = 0.91; Δ peak latency: 19 ± 45 ms, p = 0.34; APV+NBQX vs. CGP35348 Δ onset latency: 255 ± 79 ms, p = 5 x 10^-4^; Δ peak latency: 257 ± 51 ms, p = 6 x 10^-5^.

Importantly, in glomeruli with persistent (but weakened) responses after APV+NBQX application, inhalation-linked Ca^2+^ transients were significantly delayed (Fig. 3B, D), with a mean increase in onset and peak latency of 255 ± 79 ms and 257 ± 51 ms, respectively (Fig. 3G). There was no change in latencies after CGP35348 application alone (Fig. 3G; see Fig. legend for summary statistics). An explanation for this substantial delay is that, in glomeruli receiving the strongest sensory input, glutamate concentrations are eventually able to overcome APV+NBQX blockade sufficiently to trigger spike bursts in sTCs. Overall, these results demonstrate that APV+NBQX blocks postsynaptic activation in glomeruli receiving all but the strongest OSN inputs, and even in those glomeruli, substantially weakens and delays activation of monosynaptically-driven TCs.

We next used in vivo pharmacology to test the contribution of mono-versus di- or polysynaptic glutamate signaling to inhalation-linked glutamate transients onto MT cells. We focused on MT cells projecting to piriform cortex (pcMTs) as multisynaptic excitatory drive is thought to predominately impact this subpopulation (Fukunaga et al., 2012;Gire et al., 2012;Igarashi et al., 2012). We selectively expressed SF-iGluSnFR.A184V in pcMTs via retrograde viral infection by targeting virus injections to anterior piriform cortex of Tbet-Cre mice (n = 2 mice), as described previously (Rothermel et al., 2013). We have established in a recent study that odorant- and inhalation-evoked glutamate transients in pcMTs are indistinguishable from those measured from sTCs or from the general MT cell population (Moran et al., 2021). As expected, inhalation-linked glutamate transients on pcMTs were increased by CGP35348 (Fig. 4A, B), with mean increases of 35% and 90% in the two mice, and no impact on inhalation-linked glutamate dynamics (Δ onset and peak latencies, mean ± SD, paired t-tests, n = 10 ROIs pooled across 2 mice: Δ onset, 8 ± 11 ms, p = 0.07; Δ peak, 8 ± 15 ms, p = 0.2). Surprisingly, subsequent application of APV+NBQX (1 mM/0.5 mM) had no significant impact on response amplitudes (Fig. 4A-C; see Fig. 4C legend for summary statistics). iGluR blockade did cause a very small, but statistically significant, increase in the latency of the inhalation-linked glutamate transient (Δ onset latency, 13 ± 12 ms, p = 0.01; Δ peak, 34 ± 16 ms, p = 1.2 x 10^-4^; Fig. 4D); however the magnitude of this change was very small relative to the overall dynamics of the inhalation-linked transient (Fig. 4B).

**Figure 4.**
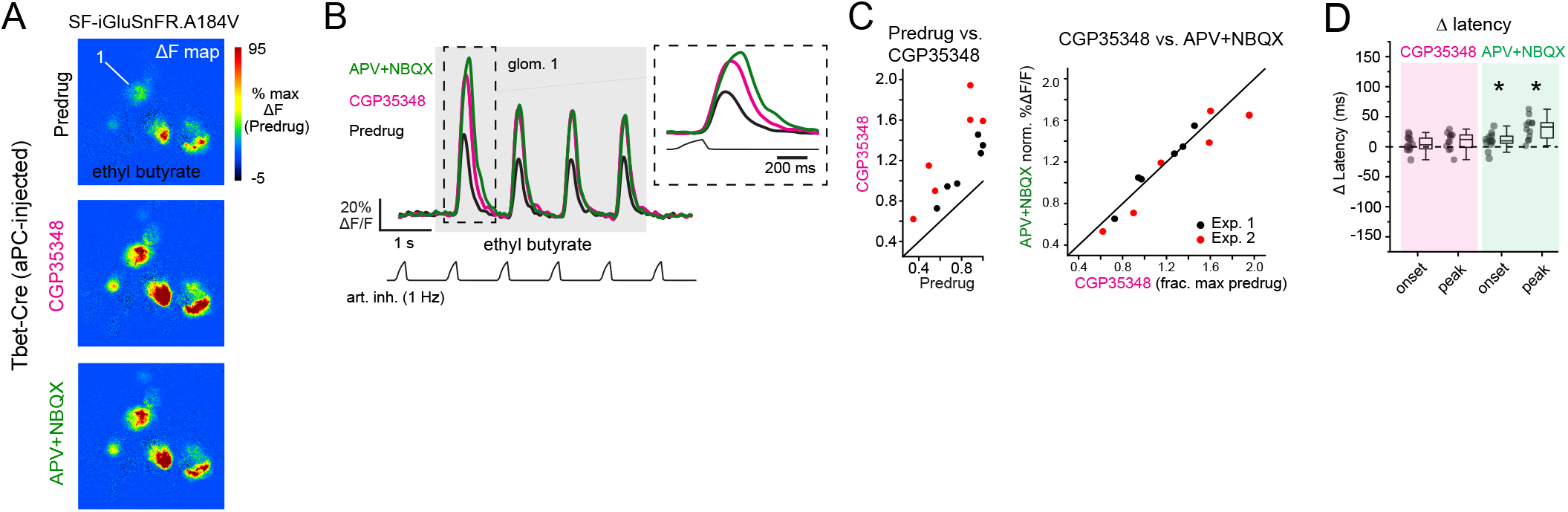
Blockade of postsynaptic excitation does not impact the magnitude or latency of inhalation-linked glutamate signaling onto piriform-projecting mitral/tufted (pcMT) cells. **A.** Odorant-evoked SF-iGluSnFR.A184V response maps across Predrug, CGP35348, and APV+NBQX conditions in an aPC-injected Tbet-Cre mouse. Maps are scaled to max of Predrug condition. **B.** Traces showing responses of glomerulus 1 (from A) across conditions. Inhalation frequency, 1 Hz. inset shows expansion of response to first inhalation of odorant in each condition, showing slight increase in latency to peak after APV+NBQX application. **C.** Comparison of peak SF-iGluSnFR.A184V responses (1^st^ inhalation) across CGP35348 (left) and APV+NBQX (right) conditions. Points indicate glomeruli from two mice (red and black). for the two mice. Statistics for each of two experiments: CGP35348/Predrug ratios (mean ± SD, t-test compared to ratio of 1): Exp. 1, 1.35 ± 0.097, p = 5 x 10^-4^, n = 6 glomeruli; Exp. 2, 1.90 ± 0.3, p = 1.6 x 10^-3^, n = 6 glomeruli; APV+NBQX/CGP35348 ratios: Exp. 1, 1.02 ± 0.07, p = 0.94; Exp. 2, 0.91 ± 0.1, p = 0.2. **D.** Little change in onset and peak latencies of SF-iGluSnFR.A184V response to first inhalation of odorant after application of CGP35348 and APV+NBQX, for same glomeruli and animals as in (C). Asterisks indicate significant change in latency after drug treatment (see Text for statistics). Latency measurements were pooled across mice due to the lower number of ROIs supporting reliable measurement due to signal-to-noise considerations.

Multisynaptic glutamatergic excitation may preferentially drive MT cell responses to weak inputs (Najac et al., 2011;Vaaga and Westbrook, 2016;Gire et al., 2019), and the odorant concentrations used in the preceding experiments were relatively high (~20 – 50 ppm; Table 1). Removal of presynaptic inhibition with CGP35348 may also enhance OSN input strength to physiologically excessive levels. Thus, we next tested the impact of glutamate receptor blockade on inhalation-linked transients in pcMT cell responses to odorants presented at 10 – 100x lower concentrations (0.1 – 5 ppm), using a higher concentration of APV+NBQX (5 mM/2.5 mM) and omitting CGP35348. In addition, to allow for greater sensitivity of glutamate detection and to focus on inhalation-linked dynamics of the glutamate signal, we used the higher-affinity SF-iGluSnFR.A184S and modified the artificial inhalation protocol to generate averaged responses to a single inhalation repeated at 0.25 Hz (Fig. 5A). This protocol yielded inhalation-triggered average waveforms that showed substantial variation in onset and peak latency as well as duration, as we have reported previously (Short and Wachowiak, 2019;Moran et al., 2021). Finally, in a subset of mice (n = 2), we used dual-color imaging to simultaneously image presynaptic glutamate transients and postsynaptic Ca^2+^ signals in the apical glomerular tufts of overlapping MT cell populations (Fig. 5B), using Tbet-Cre mice crossed with the Thy1-jRGeCO1a reporter line (Dana et al., 2018) (see Methods).

**Figure 5.**
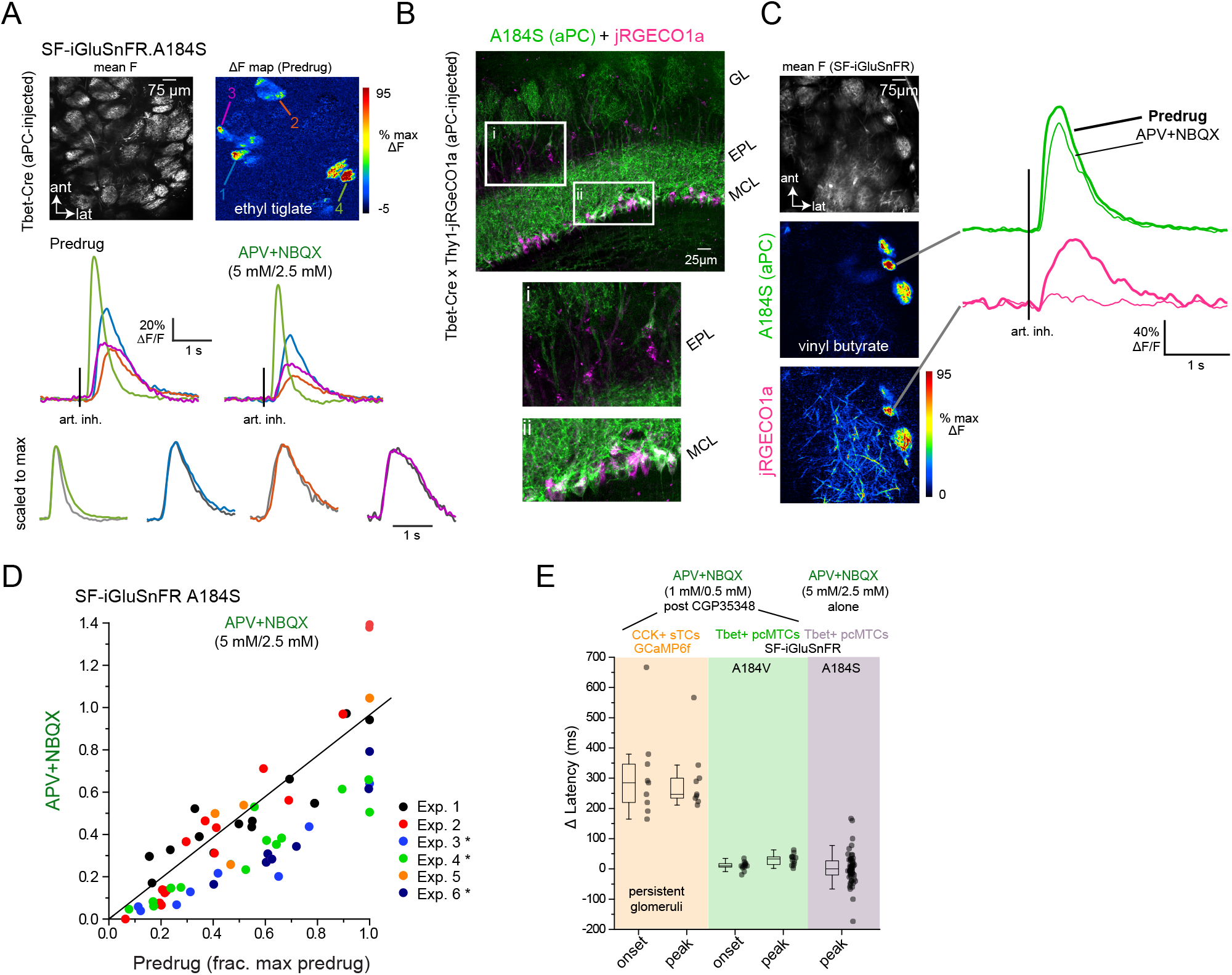
Dual-color imaging confirms small impact of multisynaptic excitation on glutamatergic input to MT cells. **A.** Top: Mean fluorescence and ΔF response maps showing odorant-evoked (ethyl tiglate, 0.7 ppm) glutamate signals in several glomeruli after SF-iGluSnFR.A184S (A148S) expression in pcMT cells. Bottom: Inhalation-triggered average (ITA) responses from four of the glomeruli, imaged before and after application of APV+NBQX (5 mM / 2.5 mM). Response amplitudes in each glomerulus are modestly reduced, but relative latencies and durations are unchanged. Lower traces show ITAs for each of four glomeruli, before and after APV+NBQX application, scaled to the same maximum (gray trace is post-drug). **B.** Dual imaging of glutamate and Ca^2+^ from overlapping subsets of MT cells. Left: Post-hoc histology showing expression of SF-iGluSnFR.A184S in pcMTs (green) and Thy1-driven jRGeCO1a expression in MT cells (magenta), in a Tbet-Cre x Thy1-jRGECO1a mouse cross. Insets highlight expression in the external plexiform layer (EPL) and mitral cell layer (MCL). Note that SF-iGluSnFR is expressed preferentially in mitral cells, while jRGECO1a is expressed in both mitral cells and tufted cells in the EPL. **C.** Left: Mean SF-iGluSnFR.A184S fluorescence and ΔF response maps for SF-iGluSnFR.A184S and jRGECO1a, imaged simultaneously in response to odorant stimulation (vinyl butyrate, 3 ppm). Right: ITA traces for the SF-iGluSnFR.A184S (green) and jRGECO1a (magenta) signals imaged from the same glomerulus, comparing Predrug (bold line) and APV+NBQX conditions. **D.** Top: ITA response amplitudes before and after APV+NBQX (5 mM / 2.5 mM) application in pcMT cells from six experiments. In each experiment, odorant was presented at two concentrations varying by a factor of 2.5 – 10; each point indicates a particular glomerulus-concentration response, before and after drug application. Responses are normalized to the maximal pre-drug response in each experiment. Asterisks indicate experiments with significant change in ITA response amplitude. Summary statistics per experiment (mean ± SD APV+NBQX/Predrug ratios, paired t-tests with Bonferroni correction, n = glom - odor concentration pairs): Exp. 1, 1,07 ± 0.35, p = 1.0, n = 13; Exp. 2, 0.83 ± 0.41, p = 1.0, n = 16; Exp. 3, 0.45 ± 0.15, p = 0.012, n = 8; Exp. 4, 0.58 ± 0.14, p = 3.7 x 10^-4^, n = 14; Exp 5, 0.97 ± 0.29, p = 1.0, n = 4; Exp 6, 0.53 ± 0.13, p = 8.2 x 10^-3^, n = 7. **E.** Little change in onset latencies and peak times of inhalation-linked responses after APV+NBQX application, measured from CCK+ GCaMP6f datasets (persistent glomeruli, same data as Fig. 3G)), pcMT SF-iGluSnFR.A184V with 1 mM/0.5 mM APV+NBQX (data in Fig. 4D)), and pcMT SF-iGluSnFR.A148S with 5 mM/2.5 mM APV+NBQX. Onset latencies were not measured for the final dataset due to the relatively low signal-to-noise ratio of these responses. Note no change in latencies for iGluSnFR signals. Boxes indicate 25^th^ and 75^th^ percentiles, line indicates median, and whiskers denote outliers with a coefficient of 1.5.

In the two dual-color imaging preparations, APV+NBQX completely blocked postsynaptic MT cell activation in 7 of 11 glomeruli analyzed, as reflected in the peak jRGECO1a Ca^2+^ signal imaged from the glomerular layer, with a reduction to 9,12,12 and 34% of pre-drug levels in the remaining 4 glomeruli. APV+NBQX had minimal impact on the glutamate transients measured in these same glomeruli (Fig. 5C). Overall, across all aPC-injected SF-iGluSnFR.A184V mice in this dataset (n = 6), the impact of APV+NBQX on the magnitude of inhalation-linked glutamate transients was modest and variable, causing a statistically significant reduction in three of six experiments (see Fig. 4 legend) and a 27 ± 26% reduction in peak ITA amplitude across all experiments (mean ± SD of mean ratios, p = 0.06, t-test, n = 6) (Fig. 5D). However, APV+NBQX did not impact the dynamics of inhalation-linked glutamate responses, with no significant change in peak times across the dataset (mean ± SD of Δ peak latencies, 9 ± 21 msec, p = 0.34, paired t-test, n = 6) (Fig. 5E). Overall, these results suggest that multisynaptic pathways modestly contribute to the strength of glutamatergic excitation of MT cells in response to low-intensity stimulation, but do not uniquely shape the inhalation-linked dynamics of this excitation.

Finally, we sought to further examine the role of direct versus multisynaptic excitation in shaping the dynamics of glutamatergic input to MT cells during repeated odorant sampling. We have shown previously that glutamate signals on MT cells show diverse temporal patterns across repeated inhalations of odorant that include adaptation, facilitation and even suppression depending on odorant identity and concentration (Moran et al., 2021). To capture this diversity, we used a 12-odorant panel (delivered concentrations ranging from 0.03 - 15 ppm) and 3 Hz artificial inhalation, comparing response patterns before and after application of APV+NBQX (1 mM/0.5 mM). This approach yielded significant odorant responses in 83 glomerulus-odorant pairs across 3 mice and included diverse excitatory response patterns (Fig. 6A). Consistent with the earlier experiments, APV+NBQX application alone had little overall impact on the magnitude of evoked glutamate signals: while there was variability in the effect for certain glomerulus-odor pairs, neither mean nor peak excitatory response showed a significant overall change after APV+NBQX application (paired t-tests, p = 0.147 for mean amplitude, p = 0.605 for peak amplitude, n = 83, Fig. 6B).

**Figure 6.**
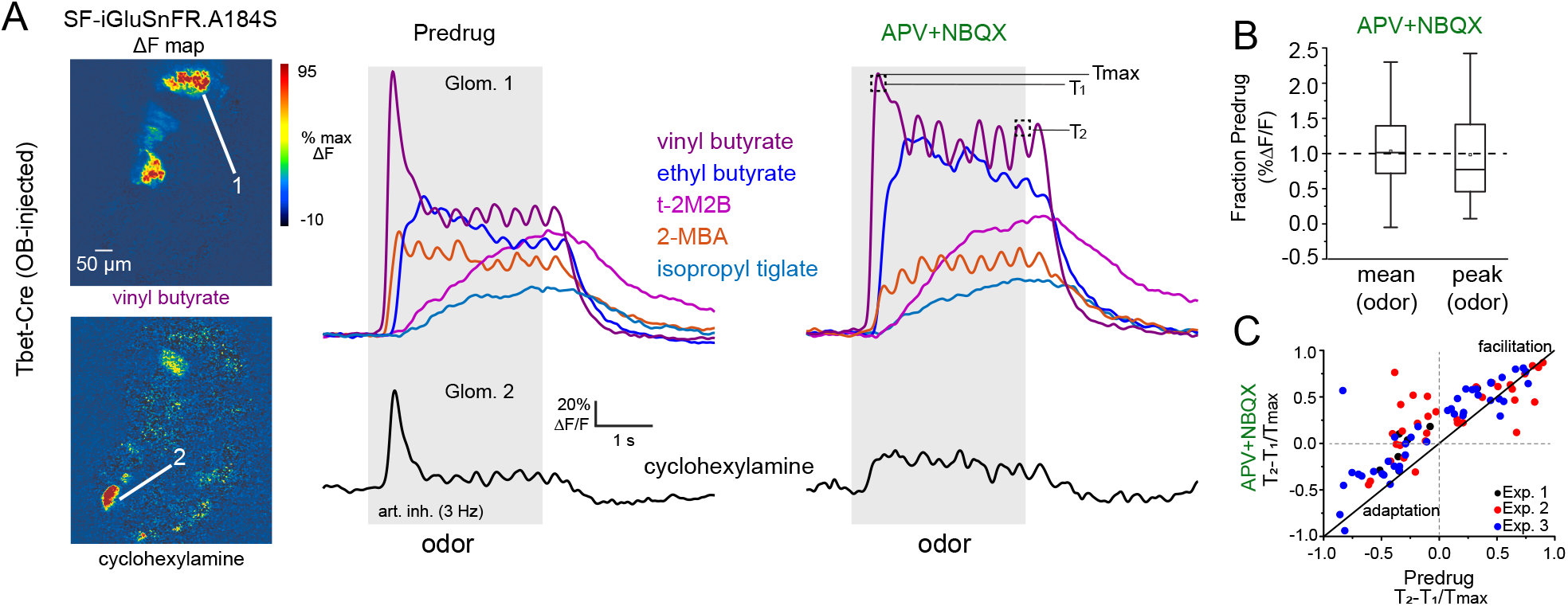
Impact of multisynaptic excitation on slow temporal dynamics of glutamate signaling across inhalations. **A.** Left: ΔF response maps showing MT cell glutamate responses to two odorants before application of APV+NBQX (1 mM/0.5 mM). Signals are from SF-iGluSnFR.A184S injected into the OB of a Tbet-Cre mouse. Right: Traces (mean of 4 presentations) from two glomeruli (top, bottom) showing diverse temporal responses to different odorants, before and after APV+NBQX application. Response to cyclohexylamine (bottom) shows a loss of the initial transient and replacement by tonic signal. Response to vinyl butyrate (top) shows increase in initial response magnitude and apparent loss of adaptation. Temporally distinct response patterns to other odorants persist after drug application. T1, T2, T_max_ indicates time-points used to quantify slow temporal dynamics, as in (C). **B.** Box plots (as in Fig. 5) showing no overall impact of APV+NBQX on mean and peak response amplitudes (each measured across the duration of odorant presentation) across all glomerulus-odorant pairs (n = 83, 3 mice). **C.** Scatter plot of T_2_-T_1_/T_max_ values before and after APV+NBQX for all responsive glomerulus-odor pairs across 3 experiments (each experiment indicated by color), showing decrease in adaptation for many pairs after application of APV+NBQX. Summary statistics: Paired t-tests, Exp. 1, n= 7 glomerulus-odor pairs, p = 0.004; Exp. 2: n = 34, p = 0.004; Exp. 3: n = 42, p = 1.63 x 10^-5^.

APV+NBQX did, however, impact the temporal patterns of the glutamate signal for a subset of responses. In particular, application of APV+NBQX alone tended to reduce adaptation that occurred following the initial inhalation of odorant (e.g., vinyl butyrate response, Fig. 6A), or it enhanced longer-lasting, tonic-type responses (e.g., ethyl butyrate response, Fig. 6A). We used a simple metric, T_2_-T_1_/T_max_, (Moran et al., 2021) to quantify and compare these adapting or facilitating dynamics before and after drug application (Fig. 6C, see also Methods). APV+NBQX caused an overall increase in T_2_-T_1_/T_max_ values (Fig. 6C), with significant increases seen in each of three preparations. These results suggest that multisynaptic pathways contribute to the slower temporal patterning of MT cell activity by modestly suppressing the excitatory drive of MT cells across repeated samples of odorant.

## Discussion

Neural circuits in the glomerular layer of the OB mediate the initial synaptic processing of olfactory inputs, yet their role in shaping the dynamic response patterns of MT cells that are thought to encode odor information during natural odor sampling remains unclear. Because of the higher complexity of MT cell response dynamics as compared to those of OSNs, including the emergence of suppressive components, much attention has focused on the role of inhibitory circuits in this process (Shao et al., 2013;Fukunaga et al., 2014;Liu et al., 2016). However, in a preceding study (Moran et al., 2021), we found that diverse temporal patterning is prominent already at the level of excitatory, glutamatergic input to MT cells in vivo, suggesting that feedforward excitatory circuits may play an underappreciated role in generating diverse MT cell responses. Here, we used *in vivo* pharmacology to more directly probe the contribution of different sources of glutamatergic input onto MT cells. Overall, our results suggest a predominant role for direct synaptic input from OSNs to MT cells in determining the dynamics of sensory-driven excitatory drive in response to odorant inhalation.

OSNs themselves show remarkable temporal diversity in their odorant responses, with activation latencies relative to inhalation spanning a range of ~250 ms, and inhalation-driven response durations varying by a similar amount in an odorant- and glomerulus-specific fashion as measured by presynaptic Ca^2+^ imaging from OSN axon terminals (Spors et al., 2006;Carey and Wachowiak, 2011;Wachowiak et al., 2013;Short and Wachowiak, 2019). While it is not surprising that this diversity persists at the level of glutamatergic signaling, our results indicate a relatively small contribution of OB circuits to further shaping inhalation-linked MT cell excitatory drive.

In addition to direct OSN inputs, feedforward, disynaptic excitation of MT cells by external tufted (ET) cells has been proposed as a primary driver of MT cell excitation (De Saint Jan et al., 2009;Gire et al., 2012). Studies from OB slices suggest that ET cells provide excitatory drive to MT cells via glutamate spillover from the ET- to - periglomerular cell synapse (Najac et al., 2011;Gire et al., 2019), allowing inhibitory circuits to regulate MT cell excitability and temporal dynamics through their inhibitory action on ET cells (Murphy et al., 2005;Gire and Schoppa, 2009;Whitesell et al., 2013;Banerjee et al., 2015;Liu et al., 2016). Here, we found that a near-complete blockade of postsynaptic activation, as confirmed with GCaMP reporters expressed in monosynaptically-driven superficial tufted cells (Sun et al., 2020), had no impact on the dynamics of inhalation-linked glutamate transients and only modestly impacted the amplitude of transients in response to low odorant concentrations. These results do not rule out some models of ET cell-mediated excitation of MT cells, which propose that the ET cell pathway is most important in regimes of weak OSN input (i.e., low odorant concentrations) (Najac et al., 2011;Vaaga and Westbrook, 2016;Gire et al., 2019), or that direct OSN inputs are shunted by gap junctions between MT cells (Gire et al., 2012), which would not be reflected in our iGluSnFR recordings. Disynaptic excitation might also provide tonic glutamatergic drive to MT cells via ET cell bursting in vivo, allowing for modulation of MT cell excitability by inhibitory circuits (Hayar et al., 2004b;Hayar and Ennis, 2007). Our findings illustrate the difficulty in extrapolating from OB slice experiments to OB circuit function in vivo; for example, glutamate spillover may contribute less to MT cell drive in vivo than in OB slices due to differences in glutamate transporter efficacy or in the dynamics of odorant-evoked glutamate release from OSN inputs.

A potential concern in interpreting these results is that the in vivo pharmacological approach was ineffective at completely blocking odorant-evoked glutamatergic transmission. While one previous study used higher concentrations of APV (50 mM) and NBQX (5 mM) to block glutamatergic transmission in vivo (Gurden et al., 2006), most prior studies have used concentrations comparable or lower than the 5 mM/2.5 mM concentrations used here to block postsynaptic activation in the OB in vivo (Pírez and Wachowiak, 2008;Fletcher et al., 2009;Lecoq et al., 2009;Shang and Xing, 2018). We also found that the 2.5 mM NBQX concentration was near the limit of its solubility in normal ringers solution, and that higher NBQX concentrations introduced severe optical interference with the imaging protocol. Nonetheless, it is conceivable that iGluR blockade failed to blocking ET cell activation, and that, in the Ca^2+^ imaging experiments used to verify effective blockade we failed to observe persistent ET cell activation due to a lack of GCaMP expression in these cells. Several lines of evidence suggest this is unlikely. First, GCaMP signals imaged from CCK-ergic sTCs, which, like ET cells, receive strong monosynaptic input from OSNs (Sun et al., 2020), were completely blocked in most glomeruli and strongly reduced and delayed in the remainder. Second, despite this delay in the persistent sTCs response, we observed no corresponding delay in the iGluSnFR response after APV+NBQX treatment. Third, in simultaneous dual-color SF-iGluSnFR and jRGECO1a imaging experiments, we saw only a modest reduction in SF-iGluSnFR response amplitude and no change in SF-iGluSnFR response latency, despite a complete or near-complete blockade of postsynaptic responses imaged in the same glomeruli. Given these results, it seems unlikely that ET cell activation and subsequent glutamate release could have been unperturbed (and undetected) by the high iGluR blocker concentrations used here.

Blocking presynaptic modulation of transmitter release from OSNs using GABA_B_ receptor antagonists impacted the magnitude, but not the dynamics, of inhalation-linked glutamate signaling onto MT cells. This result is predicted from earlier studies based on Ca^2+^ imaging from OSN terminals or synaptopHluorin-based measurements of transmitter release, which found that presynaptic inhibition mediates gain control of sensory input to the OB (McGann et al., 2005;Wachowiak et al., 2005). Blocking presynaptic inhibition had no impact on the presence of suppressive responses in MT cells, as measured with Ca^2+^ imaging, indicating that this suppression is mediated by feedforward inhibition rather than inter- or intraglomerular suppression of ongoing drive from OSNs. In some cases, GABA_B_ receptor blockade had less impact on the first inhalation of odorant than on subsequent inhalations (i.e., Fig. 1E), an effect that we have not observed with presynaptic Ca^2+^ imaging (compare Fig. 1B and Fig. 1E; see also (Pírez and Wachowiak, 2008)). This result is expected if the initial inhalation-driven burst of OSN activity is sufficiently fast and synchronous to drive glutamate release before GABA_B_-mediated inhibition can be activated, which takes approximately 50 ms to reach maximal strength due to G-protein coupled signaling in the presynaptic terminal (Wachowiak et al., 2005).

Glutamate signaling also shows substantial temporal diversity across repeated inhalations of odorant in both anesthetized and awake mice (Moran et al., 2021), and blocking multisynaptic activation with APV+NBQX altered these temporal patterns, generally by reducing adaptation of the glutamate signal or enhancing its facilitation across successive inhalations. This effect is consistent with a disinhibition of glutamatergic drive by APV+NBQX, and could reflect the removal of feedback presynaptic inhibition of OSN terminals (Wachowiak et al., 2005;Shao et al., 2009) or blockade of feedforward inhibition of ET cells; GABA_B_ receptor blockade using CGP35348 prior to APV+NBQX application, as we did with low-frequency inhalations, should distinguish these possibilities. Lastly, we did not attempt to block metabotropic glutamate receptors in these experiments, although they have been implicated in diverse contributions to MT cell and ET cell excitability, including mediating prolonged odorant-evoked MT cell responses in vivo (Dong and Ennis, 2014; Dong and Ennis, 2017; Matsumoto et al., 2009). Despite the need for further dissection of the transmitter pathways involved in glomerular processing, our results suggest that multisynaptic circuits may contribute more to slow temporal patterning of glutamatergic drive to MT cells than to shaping inhalation-linked excitatory dynamics.

MT cells show a striking correspondence between temporal patterns of glutamatergic input to their dendrites and their postsynaptic activity patterns, as measured with simultaneous Ca^2+^ imaging (Moran et al., 2021). A caveat to this similarity is the possibility that the SF-iGluSnFR signal reflects dendritic release of glutamate from MT cells themselves (Isaacson, 1999;Najac et al., 2015). However, we found that near-complete pharmacological blockade of MT cell activation enhanced, rather than suppressed, responses across repeated inhalation, suggesting that the SF-iGluSnFR signal largely reflects glutamatergic input to, rather than glutamate release from, MT cells. Taken together, these results suggest that direct glutamatergic input from OSNs provides the bulk of excitatory drive to MT cells, and that diversity in the dynamics of this input may be a primary determinant of the temporal diversity in MT cell responses that underlies odor representations at this stage. Experiments using glutamate or other transmitter reporters, multiplexed with measures of postsynaptic activation, will be important in further unraveling the contributions of OB circuits to shaping OB outputs in vivo, and as a function of odorant sampling in the behaving animal.

## MATERIALS AND METHODS

### Animals

Experiments were performed on male and female mice expressing Cre recombinase (Cre) in defined neural populations. Mouse strains used were: Pcdh21-Cre (Tg(Pcdh21-cre)BYoko), Gensat Stock #030952-UCD; OMP-Cre (Tg(Omp-tm4-Cre)Mom), JAX Stock #006668, Tbet-Cre (Tg(Tbx21-cre)1Dlc), JAX Stock #024507, CCK-IRES-Cre (Tg(CCK-IRES-Cre)Zjh), JAX Stock #012706(Haddad et al., 2013), and Thy1-jRGeCO1a Tg(Thy1-jRGECO1a)GP8.31Dkim/J, JAX Stock #030526 (Dana et al., 2018). Mice ranged from 3-8 months in age. Mice were housed up to 4/cage and kept on a 12/12 h light/dark cycle with food and water available ad libitum. All procedures were carried out following the National Institutes of Health Guide for the Care and Use of Laboratory Animals and were approved by the University of Utah Institutional Animal Care and Use Committee.

### Viral vector expression

Viral vectors were obtained from the University of Pennsylvania Vector Core (AAV1 or 5 serotype, AAV.hSynap-FLEX.iGluSnFR, AAV1 serotype AAV.hSynap-FLEX.GCaMP6f), HHMI Janelia Campus or Vigene (AAV1 or 5 serotype, pAAV.hSynap-FLEX.SF-iGluSnFR.A184V, pAAV.hSynap-FLEX.SF-iGluSnFR.A184S). Virus injection was done using pressure injections and beveled glass pipettes, as described previously (Rothermel et al., 2013;Wachowiak et al., 2013;Short and Wachowiak, 2019). After injection, mice were given carprofen (Rimadyl, S.C., 5 mg/kg; Pfizer) as an analgesic and enrofloxacin (Baytril, I.M., 3 mg/kg; Bayer) as an antibiotic immediately before and 24 hours after surgery. Mice were singly housed after surgery on ventilated racks and used 21-35 days after virus injection. In some mice, viral expression was characterized with post-hoc histology using native fluorescence.

### In vivo two photon imaging

Two-photon imaging in anesthetized mice was performed as described previously (Wachowiak et al., 2013;Economo et al., 2016). Mice were initially anesthetized with pentobarbital (50 - 90 mg/kg) then maintained under isoflurane (0.5 – 1% in O_2_) for data collection. Body temperature and heart rate were maintained at 37 °C and ~ 400 beats per minute. Mice were double tracheotomized and isoflurane was delivered passively via the tracheotomy tube without contaminating the nasal cavity (Eiting and Wachowiak, 2018). Two-photon imaging occurred after removal of the bone overlying the dorsal olfactory bulb.

Imaging was carried out with a two-photon microscope (Sutter Instruments or Neurolabware) coupled to a pulsed Ti:Sapphire laser (Mai Tai HP, Spectra-Physics; or Chameleon Ultra, Coherent) at 920-940 nm and controlled by either Scanimage (Vidrio) or Scanbox (Neurolabware) software. Imaging was performed through a 16X, 0.8 N.A. objective (Nikon) and emitted light detected with GaAsP photomultiplier tubes (Hamamatsu). Fluorescence images were acquired using unidirectional resonance scanning at 15.2 or 15.5 Hz. For dual-color imaging, a second laser (Fidelity-2; Coherent) was utilized to optimally excite jRGECO1a (at 1070nm) and emitted red fluorescence collected with a second PMT, as described previously(Short and Wachowiak, 2019;Moran et al., 2021).

### In vivo pharmacology

In vivo pharmacology was carried out after removing the bone and dura overlying the dorsal olfactory bulb, using protocols described previously (Pírez and Wachowiak, 2008;Brunert et al., 2016). Drug solutions (1mM CGP35348 and 0.5 mM NBQX / 1 mM APV) were dissolved in Ringers solution, pre-warmed on a heating block, and applied in bulk to the dorsal OB without the use of agarose or coverslip. For higher concentration experiments, we utilized 2.5 mM NBQX / 5 mM APV dissolved in Ringers immediately prior to use. We waited at least 10 minutes after drug application to allow for absorption and temperature equilibration across the tissue. For the experiments where we applied ionotropic glutamate blockers following GABA_B_ blockade, NBQX + APV was applied immediately after CGP35348 without an intervening control wash. Due to the extremely slow washout of CGP35348 in vivo, we considered CGP35348 to still be present during the subsequent NBQX + APV application.

### Odorant stimulation

In most experiments, odorants were presented as precise dilutions from saturated vapor (s.v.) in cleaned, desiccated air using a custom olfactometer under computer control, as described previously (Bozza et al., 2004;Economo et al., 2016). Odorants were presented for durations ranging from 2 - 8 sec and for single sniff, Inhalation Triggered Averages (ITA)) the odorant duration was 70 sec. Artificial inhalation rates spanned 0.25 – 2 Hz. Clean air was passed across the nostrils in between trials to avoid contribution from extraneous odorants in the environment. Odorants were pre-diluted in solvent (1:10 or 1:25 in mineral oil or medium chain triglyceride oil) to allow for lower final concentrations and then diluted to concentrations ranging from 1% to 3% s.v. Relative increases in concentration were confirmed with a photoionization detector (miniRAE Lite, PGM-7300, RAE Systems) 3 cm away from the flow dilution port. Estimated final concentrations of odorants used ranged from 0.03 – 48 ppm, depending on vapor pressure and s.v. dilution (Table 1). For experiments testing a larger panel of 12 odorants (e.g., Fig. 6), we used a novel olfactometer design that allowed for rapid switching between odorants with minimal cross-contamination (Burton et al., 2019). Here, odorants were presented for 3 seconds, in random order, using 10 second interstimulus intervals (Table 1) and repeated 3 times each. Odorant presentation was as described in a previous publication (Burton et al., 2019), using an eductor nozzle for additional mixing in a carrier stream of filtered air. The end of the eductor was placed 5 -7 cm from the nose. With the configuration used, estimated dilutions of odorant were approximately 1.5% s.v.; odorants were prediluted to achieve relatively sparse activation of dorsal glomeruli (Burton et al., 2019). Estimated final concentrations ranged from 0.03 – 15 ppm.

### Data Analysis

Image analysis was performed using custom software written in MATLAB (Mathworks). For display, odorant response maps were displayed using ΔF values rather than ΔF/F to minimize noise from nonfluorescent regions. Activity maps were scaled as indicated in the figure and were kept to their original resolution (512 x 512 pixels) and smoothed using a Gaussian kernel with σ of 1 pixel. For time-series analysis, regions of interest (ROIs) were chosen manually based on the mean fluorescence image and were further refined based on odorant response ΔF maps, then all pixels averaged within an ROI to generate time-series for analysis. Time-series were computed and displayed as ΔF/F, with F defined as the mean fluorescence in the 1-2 sec prior to odorant onset, and upsampled to 150 Hz for analysis using the MATLAB pchip function.

To analyze response patterns, traces were averaged across 4-8 presentations of each odorant. Responses were classified as having significant excitatory and/or suppressive components (excitatory for Fig. 1-6, suppressive only for Fig. 2) as follows. First, each averaged response in the Predrug condition was z-scored using a baseline from 0.5 to 2 s before odorant onset, with z defined as the SD of the baseline period concatenated for each odorant response spanning each ROI. Peak excitation was measured as 95% of the maximum signal during the duration of the odorant presentation (4 – 8 sec); when applicable, suppressive responses (i.e., Fig. 2) were measured as the 15th percentile of all values in a time window from odorant onset to 500ms after odorant offset. We then used a conservative criterion for significance of z = ± 4 SD for identifying significant excitatory or suppressive responses for further analysis.

For analysis of inhalation-linked dynamics, in most experiments, responses to the first inhalation after odorant onset were analyzed, after averaging across 4 - 8 presentations. In other experiments (e.g., Fig. 5) inhalation-triggered average (ITA) responses were generated by averaging each inhalation, repeated at 0.25 Hz, over a 70 sec odorant presentation (17 inhalations averaged in total), as described previously (Short and Wachowiak, 2019). Onset and peak latencies, relative to the start of inhalation, were calculated from the ΔF/F time-series using the ‘risetime’ function in MATLAB to fit a rising sigmoid to the baseline and peak ΔF/F levels and return the time to 10% of the rise from baseline to peak (onset latency) and the time to 90% of rise from baseline to peak (peak latency). Latency values were calculated after upsampling but with no additional temporal filtering.

For analysis of odorant-evoked dynamics across multiple sniffs (i.e., Fig. 6), responses to 4 - 8 presentations of odorant were averaged before analysis. Changes in response amplitude over time, or T_2_-T_1_/T_max_, were calculated as the difference in amplitude between the peak ΔF/F following the first (T_1_) and the second-to-last (T_2_) inhalation during odorant presentation, divided by the maximum ΔF/F during the 4 sec odor presentation.

For statistical analysis of drug effects, each experiment was treated as an independent observation, since drug treatment was applied across all ROIs in an experiment. For datasets with 5 or more experiments, summary statistics are reported as the mean (or median, for non-normally-distributed data) ± standard deviation (SD) effect across all imaged ROIs, and comparisons made on these mean/median values. For datasets with fewer than 5 experiments, mean (or median) effects across ROIs are reported for each experiment and, when possible given the number or ROIs, tests for significance performed on each experiment, using ROIs as the independent measure, and p-values adjusted for multiple comparisons based on the number of experiments per dataset. Statistical tests were performed in Origin (OriginLab Corp.). All measurement of response parameters was done using analysis code that was independent of treatment or comparison condition.

Data and analysis code underlying the results of this study are available from the corresponding author upon request.

## Author Contributions

A.K.M and M.W. designed research. A.K.M, T.P.E, and M.W. performed research, A.K.M., T.P.E., and M.W. analyzed data, A.K.M., T.P.E. and M.W. wrote the paper.

## Acknowledgments

We thank L. Looger and J. Marvin and the GENIE Project from HHMI Janelia Research Campus for sharing the SF-iGluSnFR virus. We also thank G. Vasquez-Opazo, R. Kummer, and J. Ball for technical assistance, T. Rust for data analysis software, as well as S. Burton, S. Short, I. Youngstrom, K. Podgorski, and D. Restrepo for providing helpful feedback and discussion.

